# Heterologous Expression of the carbon monoxide dehydrogenase Gene from *Clostridium* sp. to Enhance Acetic Acid and Alcohol Production from CO₂

**DOI:** 10.1101/2024.12.21.629878

**Authors:** Athmakuri Tharak, G Suresh, Sreeram Kaveti, Nishant Jain, S Venkata Mohan

## Abstract

This study evaluates the performance of carbon monoxide dehydrogenase (*codh*) embedded strains in bench-scale microbial electrochemical systems (MES) for CO₂ reduction to biofuels and biochemicals. CO_2_ fermentation efficiency was evaluated by comparing the wild-type *Clostridium acetobutylicum* (Wild), a negative control *E. coli* strain lacking the *codh* gene (NC-BL21), and engineered *E. coli* strain (Eng) alone and with IPTG induction (Eng+IPTG). Four electrochemical systems were used viz. Wild+E, NC-BL21+E, Eng+E, and Eng+IPTG+E, with a poised potential of 0.6 V applied to the working electrode. CO₂ and bicarbonate were supplemented to a total inorganic carbon (IC) concentration of 40 g/L, with a retention time of 60 h. The engineered strains demonstrated enhanced metabolic performance compared to the wild-type and negative control strains, yielding a maximum formic acid concentration of 2.1 g/L and acetic acid concentration of 7.8 g/L under the Eng+IPTG condition. Ethanol yield was highest at 3.9 g/L under the Eng+IPTG+E condition, substantially exceeding the 2.4 g/L acetic acid yield observed in the wild-type strain. The engineered strains showed superior cumulative yields (0.4075 g/g), improved *codh* charge flux stability (60 vs. 5 for Wild), and upregulated expression of genes in the Wood-Ljungdahl pathway. Bioelectrochemical performance analysis demonstrated elevated reductive catalytic currents, enhanced CO₂ reduction, and optimal charge transfer kinetics. This study highlights the effectiveness of genetic and process engineering, particularly *codh* overexpression and IPTG induction, in optimizing microbial electrosynthesis for biofuel and biochemical production from C1 gases.

## 1. Introduction

Non-genetic approaches in CO₂ and syngas fermentation, such as process optimization and metabolic regulation through environmental factors, face several limitations. Microbial gas fermentation converts CO_2_ and syngas into biofuels like acetate and ethanol, offering a sustainable alternative to fossil fuels. However, challenges such as low gas solubility and inefficient substrate utilization limit conversion efficiency. CO_2_’s low solubility (∼0.034 M in water) constrains microbial growth, while high CO and H_2_ concentrations can disrupt microbial pathways, reducing productivity (1–3). Moreover, the metabolic pathways utilized by microorganisms for gas fermentation are often inefficient, particularly in converting CO_2_ to useful products. Microorganisms such as *Clostridium* spp. and acetogens typically utilize the WLP for CO_2_ fixation, which involves complex enzymatic reactions that can be subject to regulation by various environmental factors (4, 5). The efficiency of these pathways can be further affected by the presence of competing metabolic routes, which can divert carbon flux away from desired products such as acetate and ethanol. In addition, the inherent low affinity of some microbes for gaseous substrates can hinder their overall productivity, leading to suboptimal fermentation rates (6).

To address these challenges and improve process efficiency, the overexpression of carbon monoxide dehydrogenase (*codh*) within engineered microbial systems has gained significant interest. *codh* s are a class of Ni-containing enzymes present in anaerobic carboxydotrophs, which facilitate the reversible oxidation of CO into CO_2_. These enzymes play a pivotal role in several anaerobic pathways, most notably the Wood-Ljungdahl pathway, where they contribute to CO_2_ reduction and subsequent production of acetyl-CoA (7,8). Importantly, *codh* -driven reduction of CO_2_ offers industrial promise due to its catalytic efficiency without the need for excessive overpotentials, a limitation observed in inorganic catalysts (9). Structural studies of *codh* s have revealed highly conserved architecture, typically comprising multiple iron-sulfur clusters, including a C-cluster that features a unique [Ni-Fe-S] coordination site responsible for the enzyme’s catalytic activity (10). Recent developments have focused on heterologous expression of *codh* in model organisms, such as Clostridium and Escherichia coli, with the goal of enhancing metabolic flux towards biofuel and biochemical production, including acetate and ethanol (11, 12). These engineered strains exhibit superior CO oxidation capabilities and higher yields of valuable compounds, particularly under optimized electrofermentation conditions that bolster electron supply to the microbial system.

Moreover, studies evaluating the fermentation performance of engineered strains using CO_2_ and syngas as feedstocks have demonstrated significant improvements in yield and process efficiency compared to wild-type strains (4,5, 13). These advancements suggest that *codh* overexpression, when coupled with electrochemical augmentation and metabolic engineering, holds substantial potential for the cost-effective production of biofuels and biochemicals from waste gases. In this study, the heterologous expression of *codh*, was employed in the model organisms and evaluated for the metabolic characteristics and production of acetate, showing the best CO oxidation and acetate production. Additionally, performed investigations on lac operon induction and electro-fermentation to evaluate their effects on the relative expression of the CODH gene, enzymatic activity assays, and flux stability. Metabolic and electrochemical parameters of the engineered strains were compared with wild type strain.

## 2. Materials and methods

### 2.1 Bacterial strains, plasmids and culture media

In this study, the bacterial strains, plasmids, and PCR primers utilized are detailed in Table 1.

**Table 1.**
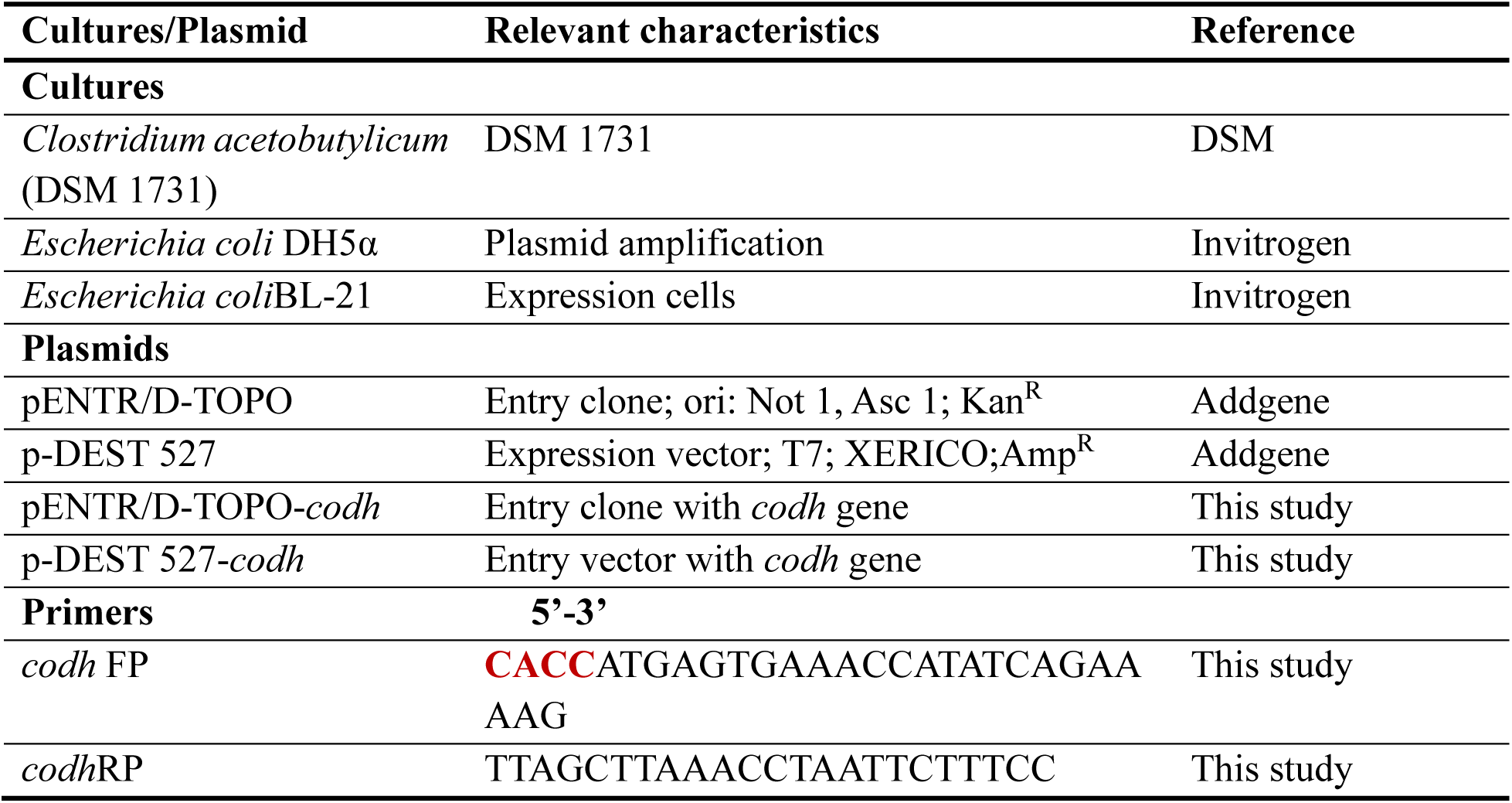
The study utilized bacterial strains, plasmids, and primers for experimental procedures.

### 2.2 Construction of the pENTR-*codh* and p-DEST- *codh* Plasmids

The codh gene ORF was amplified via PCR using primers with CACC overhangs, cloned into the pENTR/D-TOPO vector, and sequence-verified in DH5α. The pENTR-codh plasmid was used for Clonase-mediated recombination into the pDEST backbone, forming the destination vector. Recombinant *E. coli* BL21 strains expressing the codh gene were confirmed by sequencing and antibiotic selection.

### 2.3 MES Setup and Fermentation kinetics

In this experimental setup, bench-scale microbial electrochemical systems having total volume of 100 mL and a working volume of 75 mL, were used to evaluate the performance of various microbial strains under controlled conditions. The MES systems with strains included wild-type *Clostridium acetobutylicum* (Wild), *E. coli* lacking carbon monoxide dehydrogenase (NC-BL 21) considered as negative control for *codh* activity, and engineered *E. coli* strains (Eng) and Engineered strain with Isopropyl β-D-1-thiogalactopyranoside (IPTG) induction (Eng+IPTG). CO₂ along with bicarbonate (HCO_3_^-^) was supplemented intermediately (10 g/L/Day) accounting for 40 g total IC. Similar four conditions (Wild; NC-BL 21; Eng; and Eng+IPTG) were evaluated in the presence of the electrodes under regulated redox environments. Four electrochemical (E) systems denoted as wild+E; NC-BL 21+E; Eng+E; and Eng+IPTG+E. Two submerged graphite electrodes were used in microbial electrochemical systems (MES), as working and counter electrodes with active surface area of 116 cm^2^ with Ag/AgCl (3.5 MKCl) as reference electrode. To assist the CO_2_ reduction reaction near the working electrode environment a poised potential of 0.6 V was applied to the working electrode. Each strain was prepared by cultivating the in respective media and inoculated with 0.6 OD with the ratio of 10 v/v. CO_2_ gas was supplemented in head space of each reactor without disturbing biofilm activity. Retention time of 60 h maintained and analyzed for the CO_2_ fixation and conversion efficiencies. Sampling and analysis were conducted to assess the metabolic and electrochemical performance of the biocatalysts, providing insights into their efficiency and stability under different conditions.

### 2.4 Analytical Methods

Cell density was quantified using a Photopette spectrophotometer at 600 nm (OD600). Over a 24-hour period, the performance of the microbial electrochemical systems (MES) was evaluated by monitoring key metabolic parameters, including the synthesis of volatile fatty acids (VFAs) such as acetic acid, formic acid, and ethanol. Consumption of C1 gases was assessed by measuring inorganic carbon (IC) concentrations, which account for both CO₂ and bicarbonate. Metabolic product analysis, specifically for ethanol and VFAs, was conducted using a Shimadzu LC20A high-performance liquid chromatography (HPLC) system equipped with a refractive index detector (RID20A) and Rezex™ RHM-Monosaccharide H+ columns from Phenomenex (India). Prior to HPLC analysis, samples were diluted and filtered through 0.2-micron filters. The HPLC analysis was performed under isocratic conditions with an elution rate of 0.5 mL/min, using ultrapure water as the mobile phase and a sample injection volume of 20 µL.

### 2.5 Bio-electrochemical assessment

The electrochemical performance of the biocatalyst was evaluated using a potentiostat (EC-lab; Biologic VMP3) with cyclic voltammetry (CV). Cyclic voltammetry were conducted at a scan rate of 20 mV/s over a potential range from 1.0 V to −1.0 V.

### 2.6 Relative gene expressions

Microbial cells (1 mL) cultured under 100% CO₂ (1 atm) for 2 days were utilized for relative gene expression analysis of carbon monoxide dehydrogenase (*codh*). Total RNA was extracted from these cells using a kit-based protocol and immediately converted into complementary DNA (cDNA) with the TAKARA PrimeScript 1st Strand cDNA Synthesis Kit. Reverse transcription-polymerase chain reaction (RT-PCR) was performed using SYBR Green as the fluorescent probe.

### 2.7 SDS-PAGE and Western blot Analysis

Strains were grown anaerobically in liquid RCM/LB media until the optical density at 600 nm (OD600) reached approximately 2.0. At this point, 3 mL of the culture was harvested, washed with sterile phosphate-buffered saline (PBS), and then resuspended in 0.5 mL PBS. The cell suspension was subjected to ultrasonication to disrupt the cells. Following cell lysis, cell-free extracts were obtained by centrifugation. The expression of target proteins in the recombinant strains was analyzed using SDS-PAGE. Western blotting was subsequently performed with a His-tag antibody to assess the expression levels of the *codh* protein. Immunoblotting protocol was opted from our previous study (16).

### 2.8 Statistical Analysis

All experiments were conducted in triplicate unless specified otherwise. Data are presented as means with standard errors. The statistical significance of the fermentation data was assessed using t-test and two-way ANOVA having multiple variables, with a significance threshold set at p=0.05. Analysis was carried out using Graph-Pad Prism software.

## 3. Results and discussion

### 3.1 Metabolic product concentrations

#### 3.1.1 Formic acid biosynthesis

Formic acid is the primary metabolite in the WLP synthesized in the very first step after assimilation of CO_2_ by *codh* gene. However, formic acid was unstable metabolite, and it converts the succeeding folate-H4 in the WLP. The wild-type *C. acetobutylicum* did not yield formic acid throughout the experiment. This result is expected as formic acid is typically playing only a transient role in the pathway, being quickly metabolized to generate reducing equivalents or acetyl-CoA. Similar to the wild strain, NC-BL 21 did not produce formic acid due to the strain’s inability to participate in carbon dioxide reduction. However, the engineered *E. coli* strain (Eng), which overexpresses the *codh* gene, exhibited formic acid production, by maximum at 1.8 g/L by 48 h before dropping significantly to 0.4 g/L by 60 h (Fig. 1 a-b). *codh* functions bidirectionally, catalyzing both the reduction of CO_2_ to formic acid and the oxidation of CO back to CO_2_, depending on cellular redox requirements and CO_2_ availability. In the Eng system, the overexpression of *codh* likely increased CO_2_ fixation, leading to formic acid production as an intermediate metabolite in the WLP. The transient accumulation of formic acid, followed by a decline after 48 h, suggests that the strain initially produces formate from CO_2_ but then channels the formate into other metabolic pathways, converting it into downstream products such as acetyl-CoA or reducing it back to CO_2_ to maintain redox balance. IPTG induction to engineered strain (Eng+IPTG) showed higher formic acid production, reaching 2.1 g/L by 48 h followed by decline further (1.1 g/L at 60 h). The IPTG induction leads to increased expression of the *codh* gene, thus elevating the metabolic flux through the CO_2_ reduction pathway. As a result, formic acid is produced in higher quantities due to the enhanced conversion of CO_2_ to formate. However, the drop in formic acid concentration after 48 hours suggests that, similar to the Eng. condition, the strain starts to metabolize formate into downstream products.

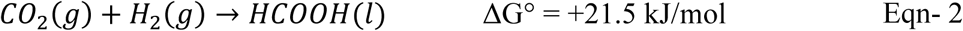

**Figure 1:**
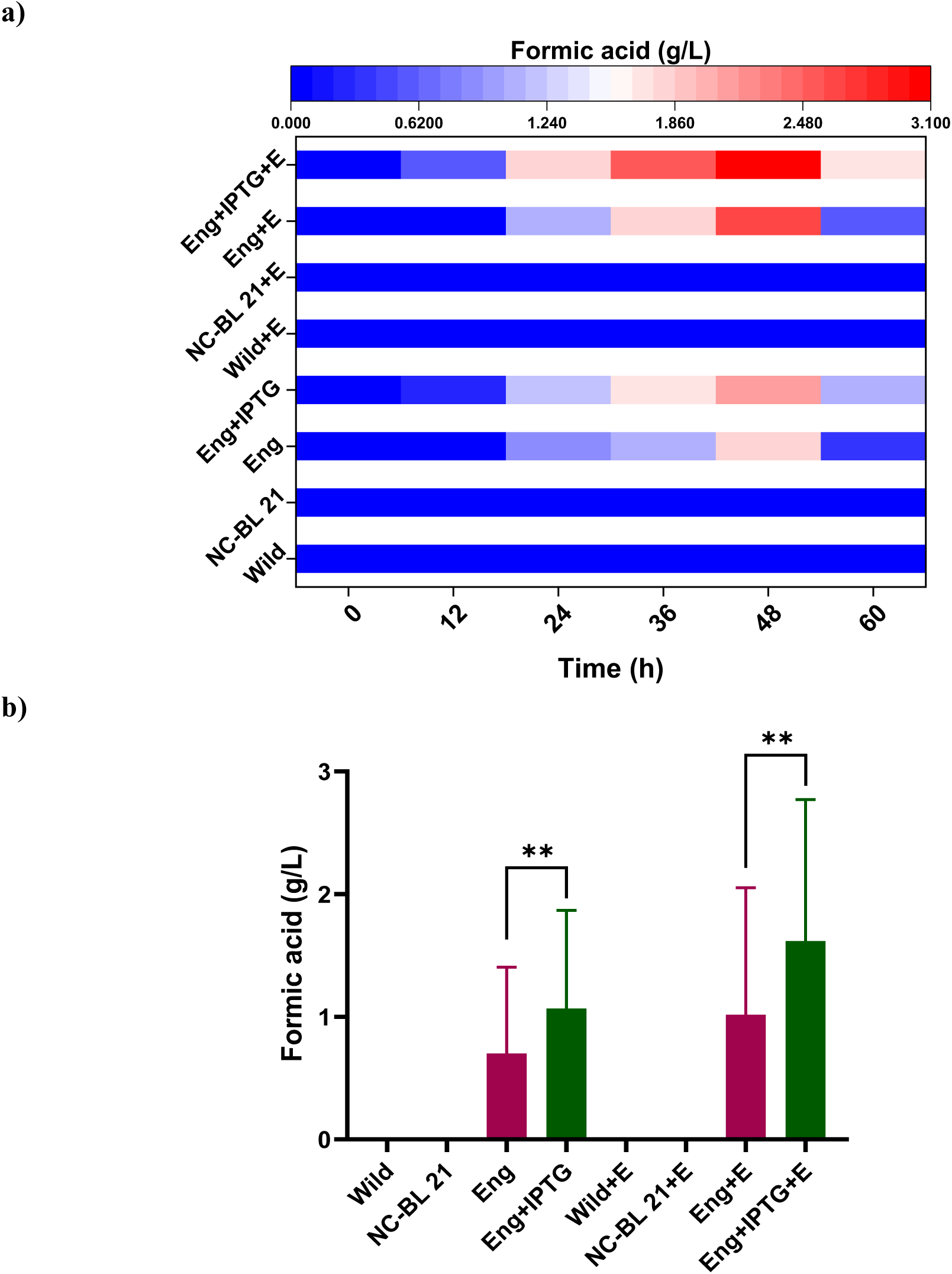
Concentration of formic acid among the systems operated, a) formic acid profile with the function of time, b) Statistically significant difference (p= 0.05 to <0.001) formic acid yield in the system operated. All p-values are based on paired t-tests corrected for multiple comparisons. Error bars are standard error of the mean (* indicates significance; ** Indicate moderate significance and ns is non-significant relation)

Wild-type *C. acetobutylicum* and NC-BL21 under electrofermentation conditions did not produce formic acid, as they lack pathways utilizing formate significantly. In contrast, engineered strains (Eng+E and Eng+IPTG+E) achieved peak formic acid yields of 2.6 g/L and 3.1 g/L at 48 h, facilitated by CODH overexpression and electrofermentation. Electrofermentation enhanced reducing equivalent availability, boosting CO_2_ reduction to formic acid, though its production declined after 48 h due to further metabolism or re-oxidation.

#### 3.1.2 Metabolic product concentrations – Acetic acid

Acetic acid production varied across MES systems, with wild-type *C. acetobutylicum* reaching 2.4 g/L at 48 h, declining to 2.1 g/L by 60 h due to metabolic limitations. Negative Control (NC-BL21) showed no acetic acid production, confirming its inability to fix CO2. The engineered strain (*Eng*) containing the *codh* gene produced significantly higher acetic acid (6.1 g/L at 48 h, 2.54-fold higher than wild-type), while IPTG induction further enhanced production to 7.8 g/L, demonstrating the impact of *codh* overexpression and induction on carbon fixation efficiency. The stoichiometric conversion of CO_2_ to acetic acid in the absence of electrode and in presence of electrode follows

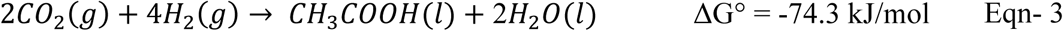

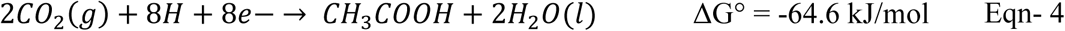

In the Wild+E condition, where *C. acetobutylicum* was subjected to electrofermentation, acetic acid production reached 2.8 g/L at 48 hours and 2.6 g/L at 60 hours, which is only a slight improvement compared to the wild-type strain without electrofermentation (Fig. 2 a-b). The relatively modest increase in productivity can be attributed to the strain’s limited ability to utilize the additional reducing equivalents provided by the electrode. Electro fermentation improves redox balance by supplying external electrons, however *C. acetobutylicum* with native WLP may not have the metabolic machinery to fully capitalize on the higher reducing equivalent flux. Nevertheless, minor improvement in acetic acid production suggests that the external electron supply facilitated higher CO_2_ reduction. No traces of acetic acid was observed in negative control.

**Figure 2:**
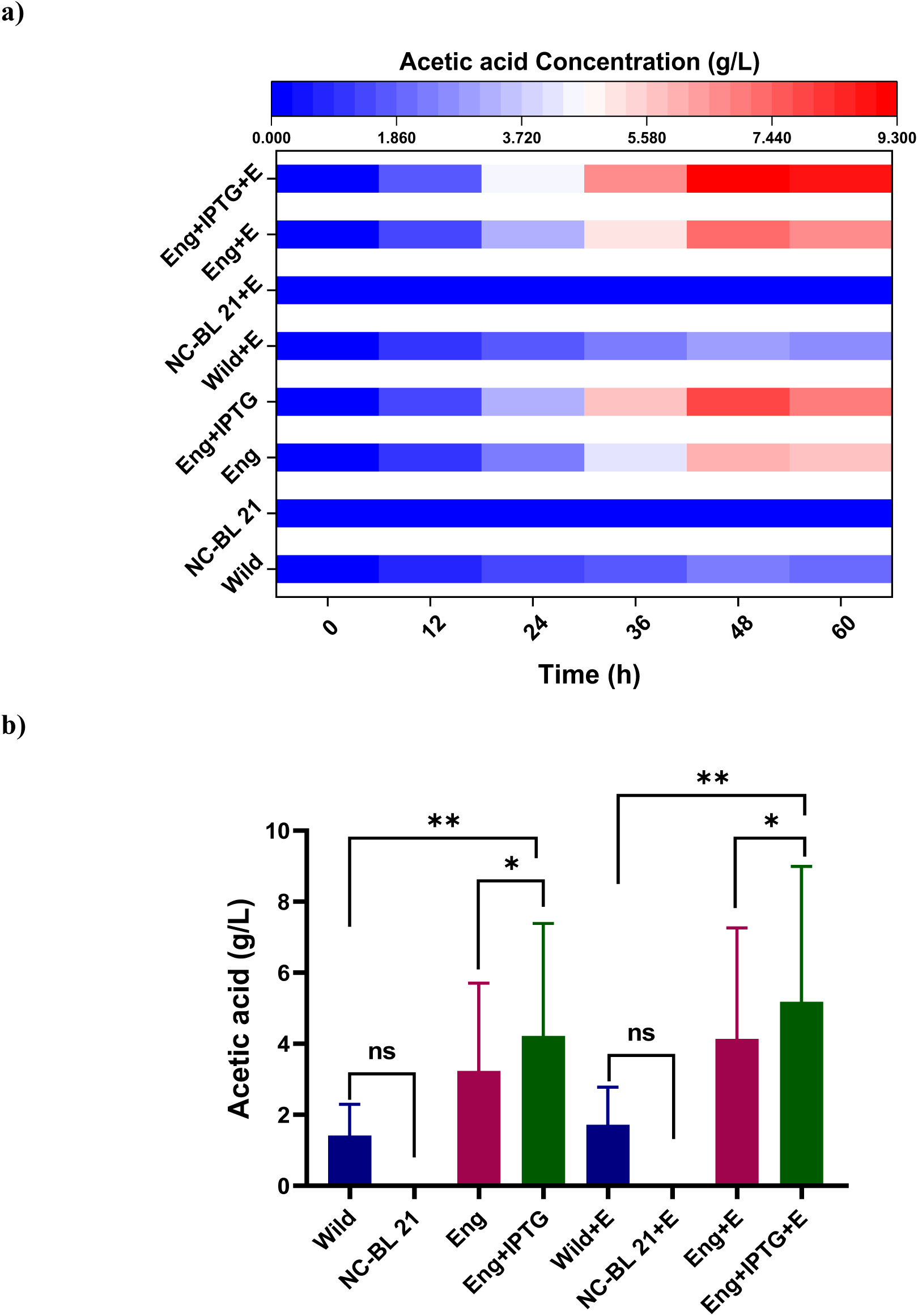
Concentration of acetic acid among the systems operated, a) Acetic acid profile with the function of time, b) Statistically significant difference (p= 0.05 to <0.001) acetic acid yield in the system operated. All p-values are based on paired t-tests corrected for multiple comparisons. Error bars are standard error of the mean (* indicates significance; ** Indicate moderate significance and ns is non-significant relation)

The electrode’s provision of reducing equivalents is insufficient to drive acetic acid production without the presence of a functional carbon fixation pathway (i.e., *codh*).

This reinforces the importance of metabolic capacity in conjunction with electrochemical inputs for acetic acid synthesis. Engineered *E. coli* with *codh* and electro fermentation (Eng+E), achieved a significant increase in acetic acid production, reaching 7.4 g/L by 48 h and 7.6 g/L by 60 h. Integration of engineered strain with electro fermentation resulted in significantly higher acetic acid titers compared to the non-electro fermentation condition (Eng). The presence of an electrode providing additional electrons increased the reducing power available for CO_2_ reduction to acetic acid. Eng+IPTG+E exhibited the highest acetic acid production among all configurations, reaching 9.3 g/L in 48 h and 8.8 g/L by 60 h. The combination of IPTG-induced overexpression of *codh* and the electro fermentation process provided an optimal environment for maximizing acetic acid synthesis. The IPTG induction amplified the metabolic flux through the CO_2_ fixation pathways, while the electrode provided an additional external electron source, further improving the overall reducing equivalent availability. This synergy between enhanced gene expression and electrofermentation likely resulted in the highest acetic acid productivity observed. The comprehensive trends indicate that *codh* overexpression, IPTG induction, and electrofermentation significantly enhance acetic acid production in *E. coli* by improving the efficiency of carbon fixation and increasing the availability of reducing equivalents. In contrast, the wild-type *C. acetobutylicum* showed modest improvement under electrofermentation due to its limited capacity to utilize external electrons. Electrofermentation proved especially beneficial when combined with genetic modifications, as seen in the Eng+E and Eng+IPTG+E conditions, highlighting the importance of metabolic engineering and electron flux management for optimizing CO_2_ conversion to acetic acid.

#### 3.1.3 Solventogenic activity – Ethanol

The ethanol production results across the eight MES systems illustrate different trends with varied strains and conditions which affect the solventogenesis. Ethanol, as a product of solventogenesis, is particularly influenced by the metabolic capabilities of the organisms, with *codh* overexpression playing a significant role in enhancing ethanol yields due to increased carbon fixation and metabolic flux towards solvent production. The wild-type *C. acetobutylicum* showed limited ethanol production, with 0.63 g/L at 48 h and dropping sharply to 0.02 g/L in 60 h (Fig. 3 a-b). Although *C. acetobutylicum* is a well-known solventogenic bacterium, typically shifting from acidogenesis to solventogenesis during later stages of fermentation under the specific conditions in current study, ethanol production remains low, likely due to the absence of significant homoacetogenic intermediates like acetyl-CoA, which are critical for solvent production. Similar to other metabolites, no traces of ethanol were observed in negative control NC-BL 21. Whereas engineered strain showed significant ethanol production, reaching 1.8 g/L at 48 h followed by decline to 0.5 g/L by 60 h. The overexpression of *codh* enhances the strain’s ability to reduce CO_2_ into formic acid and then further metabolize it into acetyl-CoA, a critical intermediate for ethanol production. Solventogenesis is triggered by the accumulation of acetyl-CoA, which gets funneled into ethanol synthesis pathways. IPTG Induction (Eng+IPTG) depicted the highest ethanol production among the non-electro fermentation systems, with 2.4 g/L at 48 h. The elevated ethanol production in this system compared to the Eng condition is due to the increased activity of the *codh* enzyme under IPTG induction. However, similar to the other conditions, the decline after 48 hours indicates that ethanol synthesis slows down as resources are depleted, or the system shifts to maintain redox balance by reducing ethanol production. Standard stichometry for the biosynthesis of ethanol follows the Eqn-5 and in case of the reduced environment of follows the Eqn-6.

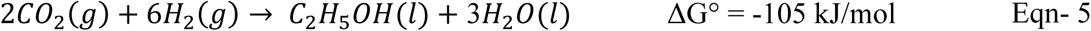

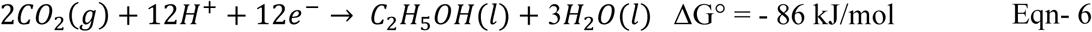

**Figure 3:**
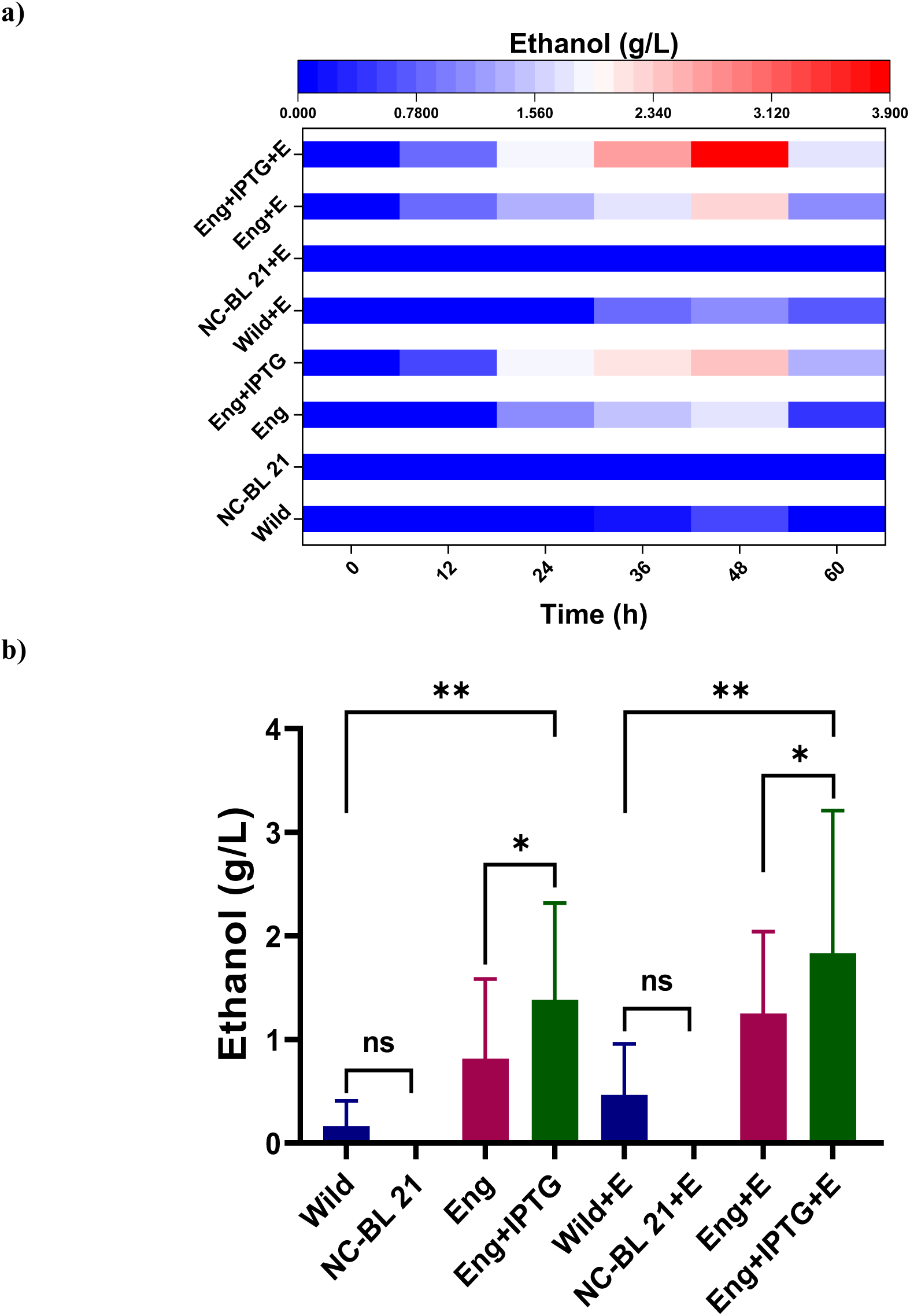
Concentration of Ethanol among the systems operated, a) Ethanol profile with the function of time, b) Statistically significant difference (p= 0.05 to <0.001) ethanol yield in the system operated. All p-values are based on paired t-tests corrected for multiple comparisons. Error bars are standard error of the mean (*indicates significance; **Indicate moderate significance and ns is non-significant relation)

Electrofermentation in the wild-type *C. acetobutylicum* (Wild+E), ethanol production increased slightly compared to the wild condition, peaking at 1.1 g/L at 48 hours. Supplies additional reducing equivalents (electrons) from the electrode, which can be used by the microbial cells to enhance the reduction of acetyl-CoA into ethanol or to acetaldehyde which is alternate metabolic route in WLP to synthesis ethanol. Due to the higher availability of reducing equivalents, which drive solventogenesis by funneling more acetyl-CoA towards ethanol synthesis. However, the improvement in ethanol production is moderate, as *C. acetobutylicum* is already naturally solventogenic, and the electrode’s impact is less pronounced than in engineered systems with greater metabolic plasticity.

#### 3.1.4 Metabolic Alterations-Effects of *codh* Recombination

*codh* catalyzes the conversion of carbon monoxide (CO) into carbon dioxide (CO₂), which feeds into the WLP. This pathway enables the production of Acetyl-CoA, which can then be converted into acetic acid, ethanol, and other metabolites. Overexpression of *codh* increases the flux of carbon through this pathway, resulting in higher production of acetic acid, ethanol, and formic acid. This explains the significantly higher g/g values for acetic acid and ethanol in the engineered strains compared to the wild type. The wild-type strain serves as the baseline, producing relatively small amounts of acetic acid and ethanol with the total metabolite yield is 0.07 g/g (Table 2). The engineered strain shows a significant improvement in metabolite production compared to the wild type. Acetic acid production increases 2.54-fold (0.15 g/g vs. 0.06 g/g in wild type), and ethanol production increases 2.86-fold (0.04 g/g vs. 0.015 g/g). The total metabolite yield is 0.24 g/g, representing a 3.2-fold increase in cumulative productivity. Upon IPTG induction, the engineered strain expresses *codh* at higher levels, resulting in further increases in metabolite production. Acetic acid production rises to 0.19 g/g, representing a 3.25-fold increase over the wild type. Ethanol production increases nearly 4-fold to 0.06 g/g, while formic acid also rises to 0.0525 g/g. The cumulative yield reaches 0.30 g/g, representing a 4.06-fold increase over the wild type. Electro-fermentation improves metabolite production in the wild-type strain. Acetic acid production increases slightly (0.07 g/g), as does ethanol production (0.02 g/g). The total cumulative yield is 0.09 g/g, reflecting only a 1.29-fold increase over the non-electro-fermentation wild type.

**Table 2:** Comparative outcomes of product yields from CO₂ by wild-type and recombinant *Clostridium acetobutylicum* strains in gas and gas-electro-fermentation systems at standard temperature and pressure (STP).

| Condition | Acetic Acid<br>(g/L) | Acetic acid<br>(g/g) | Formic Acid<br>(g/L) | Formic acid<br>(g/g) | Ethanol<br>(g/L) | Ethanol<br>(g/g) | Cumulative<br>metabolite yield (g/L) | Cumulative<br>metabolite yield (g/g) |
| --- | --- | --- | --- | --- | --- | --- | --- | --- |
| **Wild** | 2.4 | 0.006 | - | - | 0.63 | 0.015 | 3.03 | 0.075 |
| **NC-BL 21** | - | - | - | - | - | - | - | - |
| **Eng** | 6.1 | 0.152 | 1.8 | 0.045 | 1.8 | 0.045 | 9.7 | 0.24 |
| **Eng+IPTG** | 7.8 | 0.195 | 2.1 | 0.052 | 2.4 | 0.06 | 12.3 | 0.30 |
| **Wild+E** | 2.8 | 0.07 | - | - | 1.1 | 0.0275 | 3.9 | 0.09 |
| **NC-BL 21+E** | - | - | - | - | - | - | - | - |
| **Eng+E** | 7.4 | 0.18 | 2.6 | 0.065 | 2.3 | 0.0575 | 12.3 | 0.30 |
| **Eng+IPTG+E** | 9.3 | 0.23 | 3.1 | 0.0775 | 3.9 | 0.097 | 16.3 | 0.40- |

Electro-fermentation combined with *codh* overexpression in the engineered strain significantly boosts metabolite production. Acetic acid production reaches 0.185 g/g, a 3.08-fold increase over the wild type. Ethanol production also increases 3.65-fold to 0.0575 g/g. The cumulative yield matches the Eng+IPTG strain at 0.3075 g/g, representing a 4.06-fold increase in total metabolite production. Eng+IPTG+Eyields the highest productivity across all strains. Acetic acid production reaches 0.2325 g/g, a 3.88-fold increase over the wild type. Ethanol production increases to 0.0975 g/g, a 6.19-fold increase (Fig. 4a). The cumulative yield is 0.4075 g/g, representing a 5.38-fold increase in total metabolite yield over the wild type. The overexpression of *codh* in engineered *E. coli* strains significantly enhances the production of acetic acid, ethanol, and formic acid, with productivity increasing further under IPTG induction and electro-fermentation.

**Figure 4:**
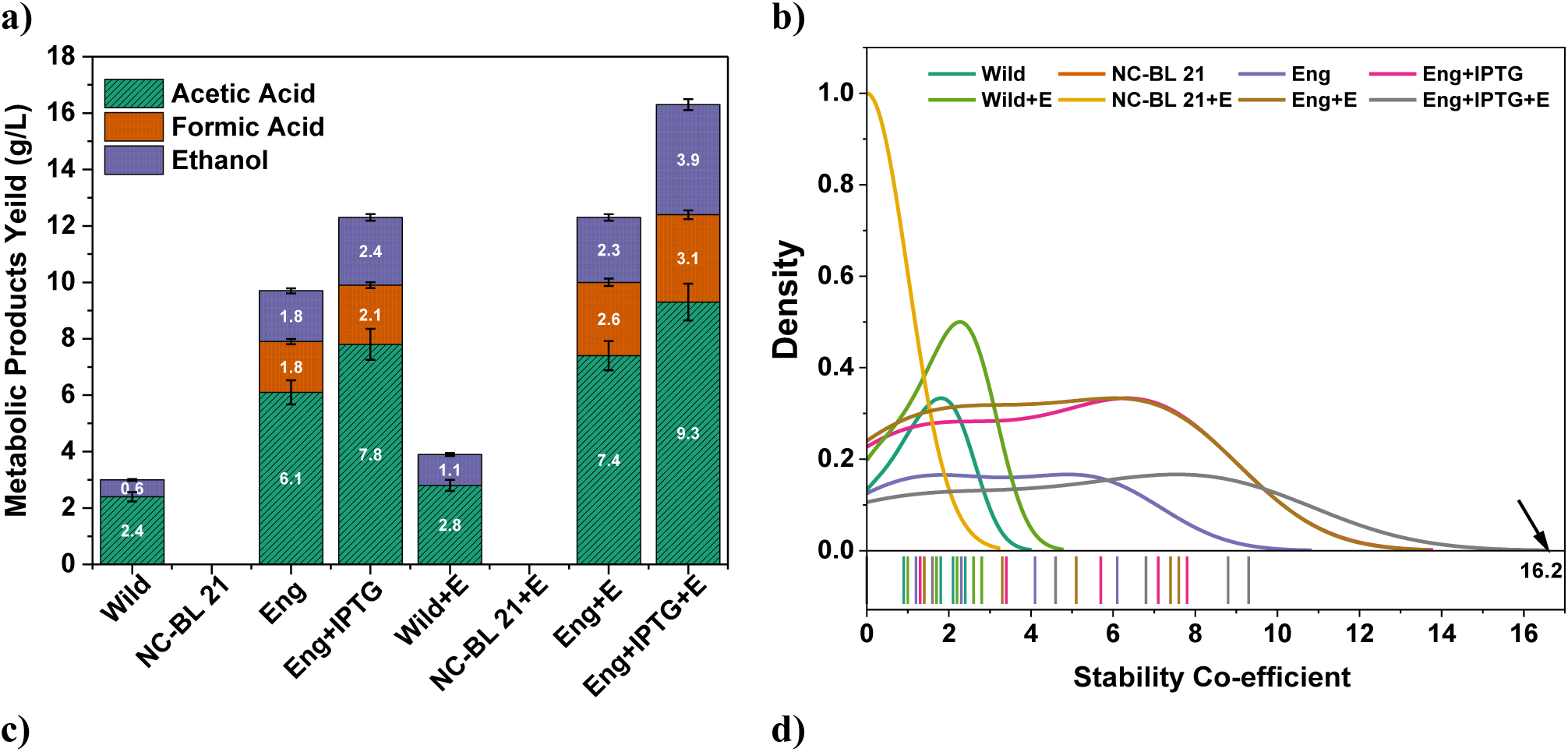
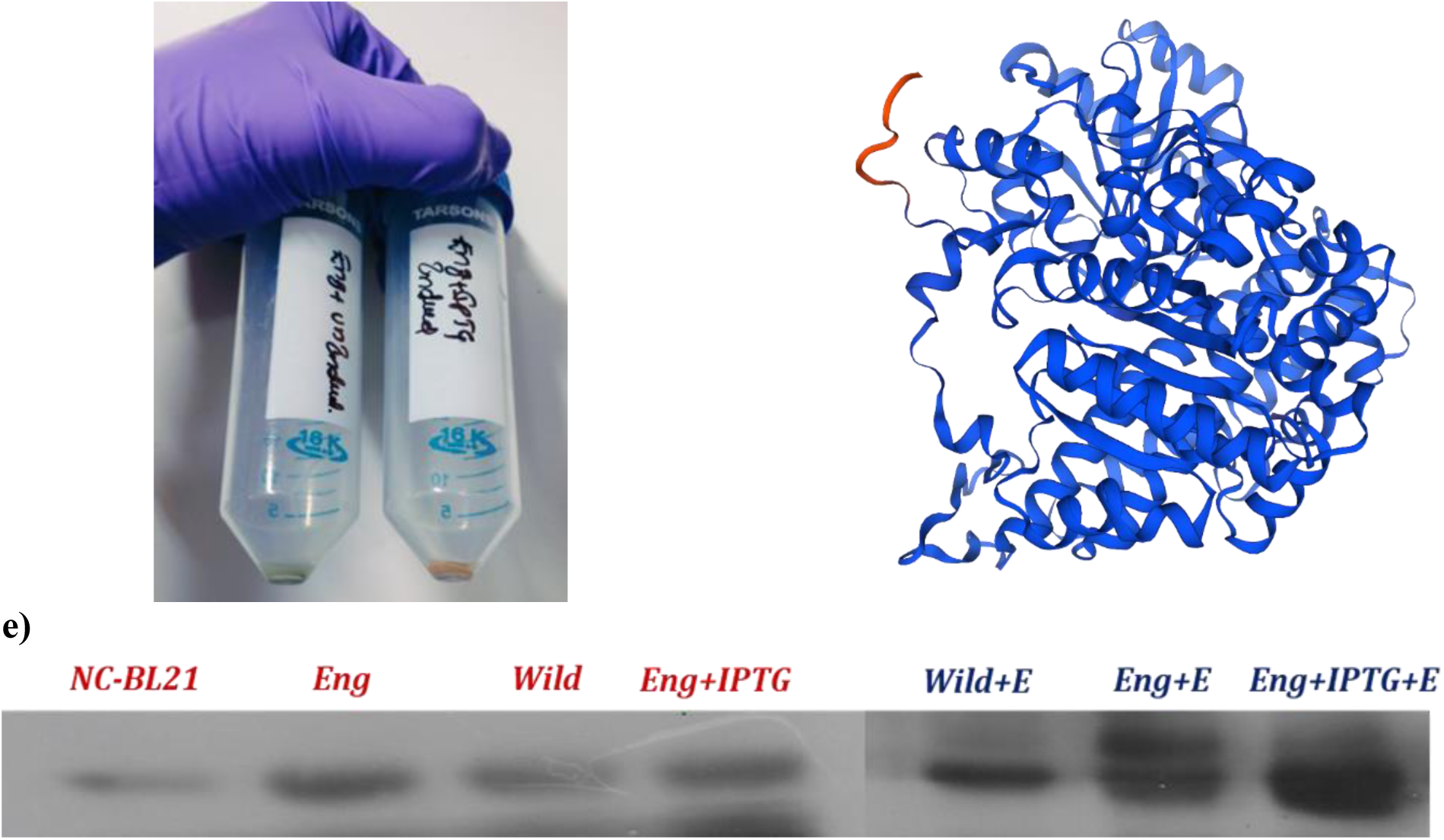
a) Cumulative metabolite biosynthesis and b) *codh* charge flux stability coefficient, c) Cell lysate of Uninduced and induced strains; d) 3D Model of the codh protein with Ni Co-factor; e) protein blot with Anti-His peroxide showing the expression levels with respect to the operation conditions.

The experimental results reveal distinct variations in *codh* flux stability across different microbial strains and conditions, with engineered strains demonstrating distinctly enhanced activity compared to the wild-type and control groups (Fig. 4b). The wild-type *Clostridium acetobutylicum* strain depicted the limited *codh* flux stability (x ≈ 5), reflecting its inherent inefficiency in electron transfer without genetic modification. The negative control strain (NC-BL 21), lacking the *codh* gene, exhibited near-zero flux stability (x ≈ 0), highlighting the critical role of *codh* in facilitating efficient electron transfer and CO₂ reduction. In contrast, the recombinant strain expressing the *codh* gene showed a significant increase in charge flux stability (x ≈ 35), highlighting *codh*’s essential role in improving electron transfer and enhancing CO₂ fixation through the WL pathway. The addition of IPTG, inducing higher *codh* expression, further increased the flux stability coefficient to approximately 50, indicating enhanced bio-electroactivity. Electro-fermentation improved the flux stability of the wild-type strain (x ≈ 15), though it remained far below the engineered strains, reaffirming the necessity of *codh* overexpression for optimal electron management in metabolite biosynthesis. Among the engineered strains, the combination of *codh* overexpression with electro-fermentation significantly enhanced stability (x ≈ 40), which was further maximized when IPTG was added (x ≈ 60). These findings clearly demonstrate that recombinant *codh* expression, particularly when induced by IPTG, significantly improves electron transfer efficiency under electro-fermentation conditions, leading to enhanced metabolic performance and increased product yields (Fig. 4c-e).

### 3.2 Relative expression of WLP genes

The expression of key genes involved in the Wood-Ljungdahl pathway was analyzed across wild-type *Clostridium acetobutylicum* (WT), engineered *E. coli* (*Eng*), and electro-fermentation (E) conditions (Fig. 5). In WT, baseline *codh* expression was 5.4, with moderate activity of associated genes. Electro-fermentation increased *codh* expression to 7.6 and upregulated related genes like *MeTr* (3.8) and *adhE1* (4.3). The engineered *E. coli* strain showed significantly higher *codh* expression (36.7), up to 66.8 with IPTG induction and electro-fermentation—a 12.4-fold increase over WT. The synergistic application of IPTG induction and electro-fermentation maximized CO2 fixation efficiency and upregulated downstream genes (*MeTr*: 18.2, *Phosphotransacetylase*: 14.6, *adhE1*: 11.3), enhancing biochemical production such as acetate and ethanol.

**Figure 5:**
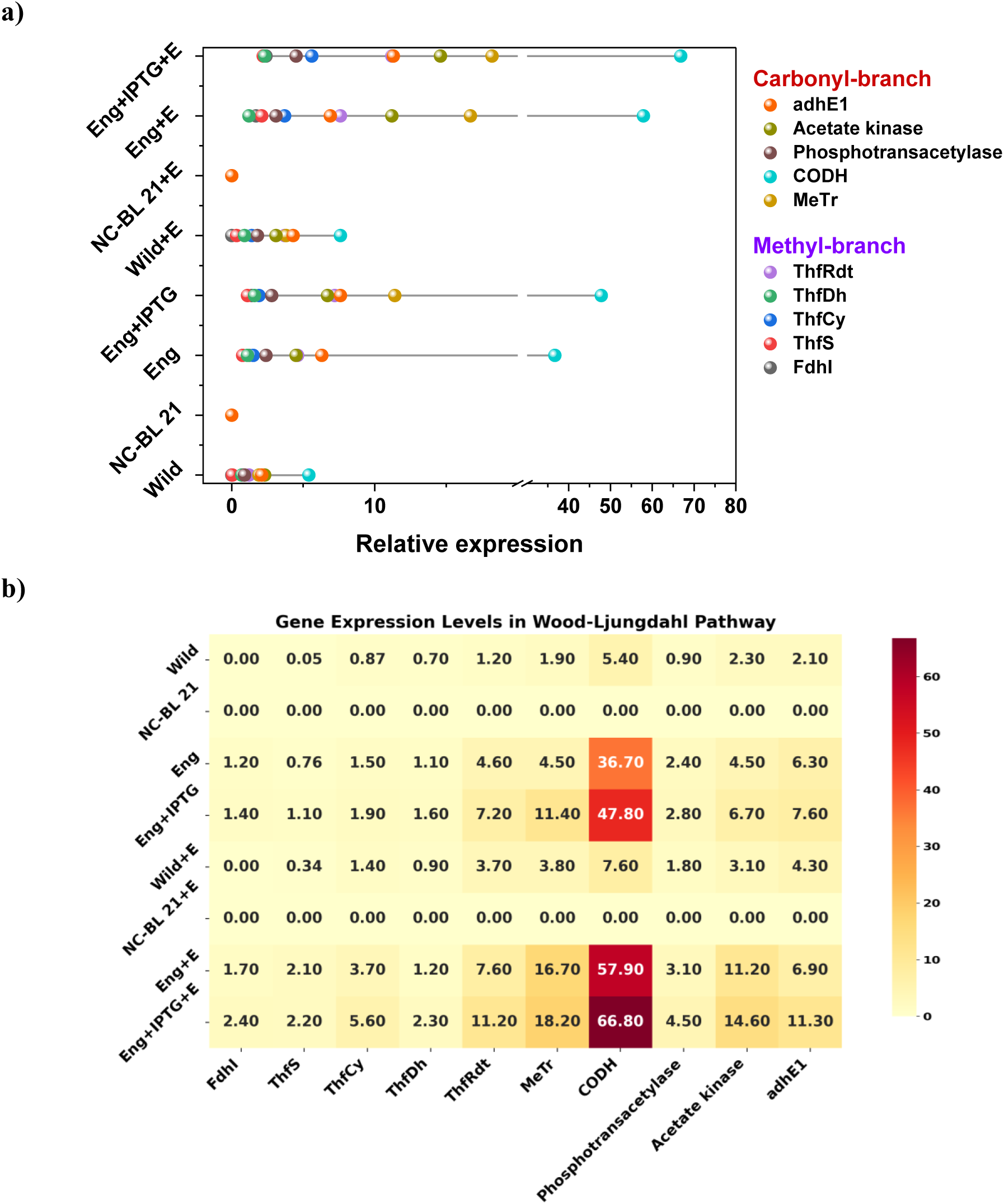
a) Relative expression of the carbonyl and methyl branch coding genes of WLP across the experimental conditions b) Heat map depicting the intensities of expression levels

### 3.4 Carbon monoxide dehydrogenase activity

The results demonstrate that carbon monoxide dehydrogenase (*codh*) activity varies significantly across the different microbial systems, reflecting the impact of *codh* overexpression, IPTG induction, and electro fermentation. The wild-type *Clostridium acetobutylicum* (Wild) shows low baseline *codh* activity (0.07 U/mg), which is slightly increased in the presence of an electrode (Wild+E, 0.09 U/mg) due to the additional reducing equivalents (Fig. 6 a-b). The negative control *E. coli* (NC-BL 21) exhibited no *codh* activity (0 U/mg), confirming its inability to reduce CO_2_ without the *codh* gene. In contrast, the engineered *E. coli* strain expressing *codh* (Eng) shows significantly higher activity (0.18 U/mg), which increases with IPTG induction (Eng+IPTG, 0.26 U/mg), indicating enhanced *codh* expression. Electro fermentation further boosts *codh* activity in engineered strains, with Eng+E showing 0.22 U/mg and Eng+IPTG+E exhibiting the highest activity (0.31 U/mg). The combination of genetic engineering, IPTG induction, and electro fermentation maximizes *codh* activity, enhancing CO_2_ reduction and carbon fixation via the Wood-Ljungdahl pathway. The external electrons provided by electro fermentation facilitate greater electron flow towards the enzymatic reduction of CO_2_ to CO, improving the conversion of inorganic carbon into organic products like acetic acid and ethanol, as previously observed in the improved CEIC and ICFR values. These findings emphasize the synergistic effects of *codh* overexpression and electro fermentation in enhancing microbial electrosynthesis efficiency.

**Figure 6:**
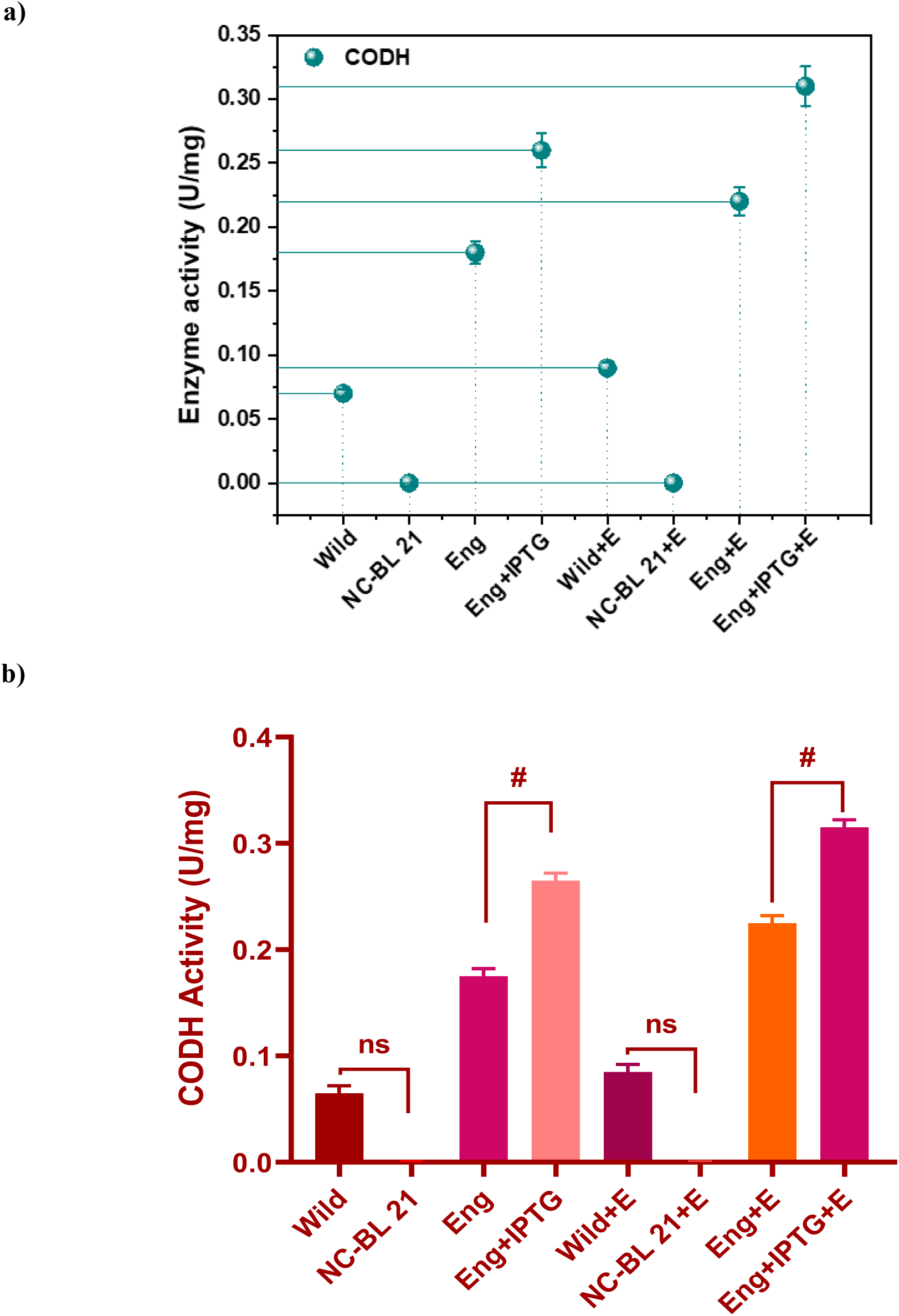
a) Specific enzyme activities of *codh* in wild-type and recombinant strains, b) Each assay was carried out in triplicates with mean and standard deviation reported.

The *codh* enzyme, a key component of the WLP, catalyzes the reversible reduction of CO_2_ to CO, playing a critical role in the autotrophic metabolism of acetogens. The nickel-iron-sulfur clusters in *codh* facilitate the transfer of electrons from reduced cofactors like MV, which was measured spectrophotometrically in this study. Higher *codh* activity correlates with improved carbon fixation, as the enzyme enhances the flow of carbon into the WLP, ultimately increasing the production of acetyl-CoA and downstream metabolites. The results align with previous studies on *Clostridium thermocaceticum*, where purified *codh* activity was found to be critical for efficient CO_2_ utilization under autotrophic conditions. The increase in *codh* activity in engineered strains, particularly when combined with electro fermentation, demonstrates the effectiveness of genetic and process engineering in enhancing microbial electrosynthesis performance. Electro fermentation, by providing an abundant supply of reducing equivalents, further stimulates *codh* activity, allowing for more efficient CO_2_ conversion and organic product formation.

### 3.5 Bio-catalyzed electrochemical reactions

#### 3.5.1 Redox currents generations

Cyclic voltammetry analysis reveals significant differences in the electrochemical behavior of the various samples. The Wild+E sample exhibits a relatively modest current response, while the NC-BL 21+E sample shows negligible current across the potential range, indicating minimal electrochemical activity. In contrast, the engineered strain (Eng+E) demonstrates a much larger current response, with distinct oxidation and reduction peaks, indicating active redox processes. The engineered strain with IPTG induction (Eng+IPTG+E) shows the highest current response, with well-defined peaks for both oxidation and reduction. The electrochemical outcomes indicate a strong correlation between the magnitude of the reduction current and CO₂ reduction activity at the working electrode. The Eng+IPTG+E condition, which exhibits the higher reductive catalytic currents (RCC) (−75 mA), demonstrates the efficient CO₂ reduction activity. This enhanced activity is likely attributed to the induced expression of *codh* gene that promote electron transfer and stimulate biocatalytic pathways for CO₂ reduction (Fig. 7a). In the Eng+E condition, the moderately high reduction current (−50 mA) suggests that genetic engineering without inducer still improves CO₂ reduction efficiency, albeit to a lesser extent. The Wild+E condition (−20 mA) represents the baseline CO₂ reduction activity of the wild strain, showing minimal electrical signals. In contrast, the NC-BL 21+E condition (−1 mA), with the least RCC, depicts the negligible reduction activity, likely due to the absence of key biocatalytic functions necessary for CO₂ reduction. The progressively increasing negative reduction currents across the conditions reflect more active electron uptake and enhanced CO₂ reduction, with the engineered strains clearly surpassing the other conditions in catalyzing CO_2_ to chemicals and fuels. Electroactive peak analysis reveals the peak positions for oxidation and reduction currents for each condition, with voltages converted to the standard hydrogen electrode (SHE) scale: Wild+E (Oxidation: +0.397 V, Reduction: −0.603 V), NC-BL 21+E (Oxidation: +0.297 V, Reduction: −0.503 V), Eng+E (Oxidation: +0.497 V, Reduction: −0.703 V), and Eng+IPGT+E (Oxidation: +0.597 V, Reduction: −0.803 V). These potentials correlate with the biological redox ladder, indicating possible involvement of electroactive compounds such as formate and NADH/NAD+ in the bio electrochemical conversion of CO_2_. The reduction peaks around −0.6 to −0.8 V potentially involving CO_2_/formate and CO_2_/acetate redox couple and oxidation peaks around +0.3 to +0.6 V involving NADH/NAD+ components (25, 26).

**Figure 7:**
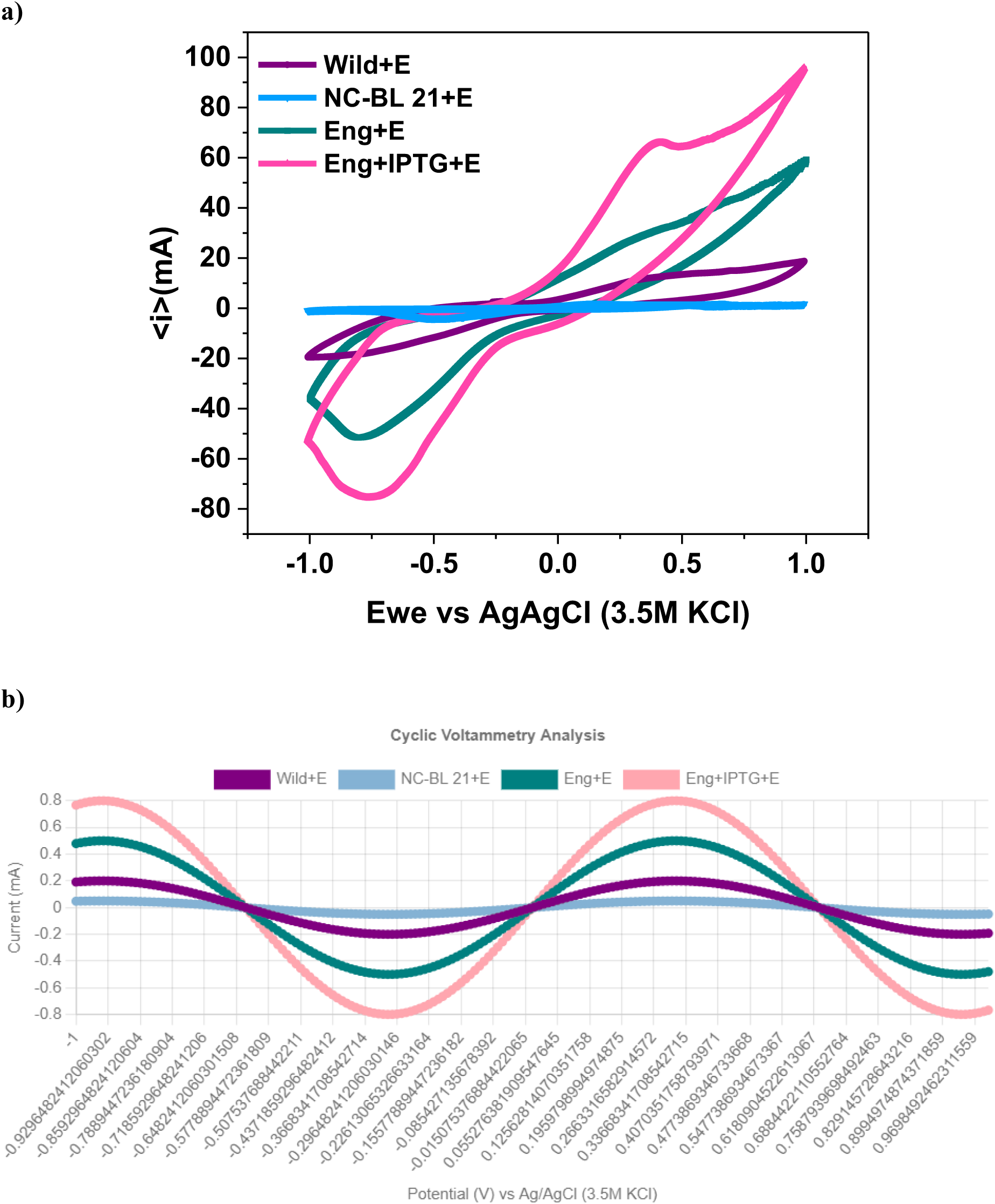
a) Cyclic voltammograms representing the electroactive species involved during the CO_2_ bio electro conversion, b) Oxidative and reductive peak analysis depicting the electrical intensities and positions.

These variations in peak currents and potentials suggest enhanced electrochemical activity in the engineered strains, likely due to genetic modifications and induced protein expression. The pronounced peaks in the Eng+IPTG+E sample indicate more efficient, possibly reversible or quasi-reversible electron transfer processes, likely facilitated by redox-active proteins, such as overexpressed *codh*. Quantitatively, the peak currents recorded for Wild+E, NC-BL 21+E, Eng+E, and Eng+IPTG+E are 0.20 mA, 0.05 mA, 0.50 mA, and 0.80 mA, respectively (Fig. 7b). The relative increases in current300% for Wild+E, 900% for Eng+E, and 1500% for Eng+IPTG+E compared to NC-BL 21+E underscore the significant enhancement of electrochemical activity in the engineered strains, particularly under IPTG induction. These results suggest that the combination of genetic modifications and IPTG induction enhances electron transfer efficiency, likely through increased biofilm formation and *codh* overexpression, which may contribute to higher product synthesis in bio-electrocatalytic systems.

## 4. Conclusion

This study investigates the metabolic product profiles and efficiencies of engineered strains relative to wild-type and control strains. Engineered *E. coli* strains exhibited a significant enhancement in formic acid production, achieving a peak concentration of 2.1 g/L at 48 hours under the Eng+IPTG condition. Acetic acid production also reached 7.8 g/L at this time point, substantially exceeding the 2.4 g/L observed in the wild-type strain. Notably, ethanol production was highest in the Eng+IPTG+E condition, yielding 3.9 g/L. The engineered strains demonstrated superior cumulative metabolite yields, with the Eng+IPTG+E condition achieving a total yield of 0.4075 g/g, alongside improved *codh* charge flux stability, evidenced by a coefficient of approximately 60, in contrast to a mere 5 for the wild-type strain. The relative expression of key genes within the Wood-Ljungdahl pathway was significantly upregulated in the engineered strains, particularly under IPTG induction and electrofermentation conditions, with *codh* enzyme activity reaching 0.31 U/mg in the Eng+IPTG+E condition, compared to 0.07 U/mg in the wild-type strain. Bioelectrochemical performance analyses indicated elevated reductive catalytic currents in the engineered strains, especially in the Eng+IPTG+E condition, suggesting enhanced electrochemical activity and CO_2_ reduction capabilities. Tafel plot analysis demonstrated improved charge transfer kinetics and biocompatibility in the engineered strains, with differential pulse voltammetry revealing peak potentials closely aligned with optimal redox potentials for efficient CO_2_ reduction. These findings underscore that the overexpression of the *codh* gene, combined with IPTG induction and electrofermentation, significantly enhances the production of acetic acid, formic acid, and ethanol in engineered *E. coli* strains. This approach improves CO_2_ fixation and conversion efficiencies, increases *codh* enzyme activity, and enhances bioelectrochemical performance, highlighting the efficacy of genetic and process engineering in optimizing microbial electrosynthesis for biofuel and biochemical production from C1 gases.

## Acknowledgements

The authors express gratitude to The Director, CSIR-IICT, for generously providing the facilities necessary to conduct this study. CSIR-IICT Communication number: IICT/Pubs./2024/444.

## References

1. Tharak A, and Venkata Mohan, 2024. Electrode-Assisted Pressurized CO Fermentation for Acetic Acid and Ethanol Production: Enhanced Carbon Fixation, Metabolic Efficiency, and Sustainability in Carbon-Negative Bioprocesses. ACS Sustainable Chemistry & Engineering. Manuscript No.: sc-2024-075377 (acssuschemeng.4c07537)

2. Lee, J., Yun, H., Feist, A.M., Palsson, B.Ø. and Lee, S.Y., 2008. Genome-scale reconstruction and in silico analysis of the Clostridium acetobutylicum ATCC 824 metabolic network. Applied microbiology and biotechnology, 80, pp.849–862.

3. Zhao, R., Liu, Y., Zhang, H., Chai, C., Wang, J., Jiang, W. and Gu, Y., 2019. CRISPR-Cas12a-mediated gene deletion and regulation in Clostridium ljungdahlii and its application in carbon flux redirection in synthesis gas fermentation. ACS Synthetic Biology, 8(10), pp.2270–2279.

4. Jones, S.W., Fast, A.G., Carlson, E.D., Wiedel, C.A., Au, J., 2016. CO_2_ fixation by anaerobic nonphotosynthetic mixotrophy for improved carbon conversion. Nat. Comm. 7, 12800.

5. Marcellin, E., Behrendorff, J.B., Nagaraju, S., DeTissera, S., Segovia, S., Palfreyman, R.W., Daniell, J., Licona-Cassani, C., Quek, L.E., Speight, R., Hodson, M.P., Simpson, S.D., Mitchell, W.P., Köpke, M., Nielsen, L.K., 2016. Low carbon fuels and commodity chemicals from waste gases-systematic approach to understand energy metabolism in a model acetogen. Green Chem. 18, 3020–3028.

6. Yu, Y., Pi, S., Ke, T., Zhou, B., Chao, W., Yang, Y., Li, Z., Li, G., Ren, N., Gao, X. and Lu, L., 2023. Artificial Soil-Like Material Enhances CO2 Bio-Valorization into Chemicals in Gas Fermentation. ACS Applied Materials & Interfaces, 15(46), pp.53488–53497.

7. Jia, D., He, M., Tian, Y., Shen, S., Zhu, X., Wang, Y., Zhuang, Y., Jiang, W. and Gu, Y., 2021. Metabolic engineering of gas-fermenting Clostridium ljungdahlii for efficient co-production of isopropanol, 3-hydroxybutyrate, and ethanol. ACS Synthetic Biology, 10(10), pp.2628–2638.

8. Nisar, A., Khan, S., Hameed, M., Nisar, A., Ahmad, H. and Mehmood, S.A., 2021. Bio-conversion of CO2 into biofuels and other value-added chemicals via metabolic engineering. Microbiological research, 251, p.126813.

9. Maru, B.T., Munasinghe, P.C., Gilary, H., Jones, S.W., Tracy, B.P., 2018. Fixation of CO_2_ and CO on a diverse range of carbohydrates using anaerobic, nonphotosynthetic mixotrophy. FEMS Microbiol. Lett. 365, fny039

10. Ragsdale, S. W. (2008). “Catalytic mechanisms of the carbon monoxide dehydrogenase/acetyl-CoA synthase complex.” Current Opinion in Chemical Biology, 12(2), 158–165.

11. Kang, H., Park, B., Oh, S., Pathiraja, D., Kim, J.Y., Jung, S., Jeong, J., Cha, M., Park, Z.Y., Choi, I.G. and Chang, I.S., 2021. Metabolism perturbation caused by the overexpression of carbon monoxide dehydrogenase/acetyl-CoA synthase gene complex accelerated gas to acetate conversion rate of Eubacterium limosum KIST612. Bioresource technology, 341, p.125879.

12. Tharak, A. and Mohan, S.V., 2021. Electrotrophy of biocathodes regulates microbial-electro-catalyzation of CO2 to fatty acids in single chambered system. Bioresource Technology, 320, p.124272.

13. Bian, B., Bajracharya, S., Xu, J., Pant, D. and Saikaly, P.E., 2020. Microbial electrosynthesis from CO_2_: Challenges, opportunities and perspectives in the context of circular bioeconomy. Bioresource technology, 302, p.122863.

14. Drake, H.L., Hu, S.I. and Wood, H.G., 1980. Purification of carbon monoxide dehydrogenase, a nickel enzyme from Clostridium thermocaceticum. Journal of Biological Chemistry, 255(15), pp.7174–7180.

15. Tharak, A. and Mohan, S.V., 2022. Syngas fermentation to acetate and ethanol with adaptative electroactive Carboxydotrophs in single chambered microbial electrochemical system. Micromachines, 13(7), p.980.

16. Bhattacharjee, D., Kaveti, S. and Jain, N., 2023. APC/C CDH1 ubiquitinates STAT3 in mitosis. The International Journal of Biochemistry & Cell Biology, 154, p.106333.

17. Alpdagtas S, Turunen O, Valjakka J, et al. The challenges of using NAD(þ)-dependent formate dehydrogenases for CO_2_ conversion. Crit Rev Biotechnol. 2021;42(6):953–972.

18. Kopke M, Held C, Hujer S, et al. Clostridium ljungdahlii represents a microbial production platform based on syngas. Proc Natl Acad Sci U S A. 2010;107(29): 13087–13092.

19. Jang, Y.S., Lee, J.Y., Lee, J., Park, J.H., Im, J.A., Eom, M.H., Lee, J., Lee, S.H., Song, H., Cho, J.H. and Seung, D.Y., 2012. Enhanced butanol production obtained by reinforcing the direct butanol-forming route in Clostridium acetobutylicum. MBio, 3(5), pp.10–1128.

20. Schmidt, M. and Weuster-Botz, D., 2012. Reaction engineering studies of acetone-butanol-ethanol fermentation with Clostridium acetobutylicum (Vol. 7, No. 5, pp. 656–661). Weinheim: WILEY-VCH Verlag.

21. Lehmann, D., Hönicke, D., Ehrenreich, A., Schmidt, M., Weuster-Botz, D., Bahl, H. and Lütke-Eversloh, T., 2012. Modifying the product pattern of Clostridium acetobutylicum: physiological effects of disrupting the acetate and acetone formation pathways. Applied microbiology and biotechnology, 94, pp.743–754.

22. Wang, J., Yang, X., Chen, C.C., Yang, S.T., 2014. Engineering clostridia for butanol production from biorenewable resources: from cells to process integration. Curr. Opin. Chem. Eng. 6, 43–54.

23. Oelgeschläger, E. and Rother, M., 2008. Carbon monoxide-dependent energy metabolism in anaerobic bacteria and archaea. Archives of microbiology, 190, pp.257–269.

24. Andersch, W., Bahl, H. and Gottschalk, G., 1983. Level of enzymes involved in acetate, butyrate, acetone and butanol formation by Clostridium acetobutylicum. European journal of applied microbiology and biotechnology, 18, pp.327–332.

25. Kumar, A., Aeshala, L.M. and Palai, T., 2024. Bioelectrochemical reduction of CO2 into formic acid using Escherichia coli whole-cell biocatalyst. Journal of Applied Electrochemistry, pp.1–12.

26. Bañeras, L., Cabeza, Á., Perona-Vico, E., Lopez-Abelarias, M., Puig, S. and De Wever, H., 2024. Microbial models for biocathodic electrochemical CO2 transformation: A comprehensive review on pure cultures. Bioresource Technology Reports, p.101766.

